# Observations on the curvature of *Physcomitrium patens* (Hedw.) Mitt. and *Funaria hygrometrica* (Hedw.) caulonemal filaments

**DOI:** 10.1101/2023.10.30.564635

**Authors:** Nick A Antonishyn, Jeffrey G Duckett, Neil W Ashton

## Abstract

Light-grown, whole gametophytic colonies of *Physcomitrium* (formerly *Physcomitrella*) *patens* (Hedw.) Mitt. exhibit a spiral morphology resulting from the strongly coordinated curvature of the population of peripheral caulonemal filaments. The direction of curvature is predominantly clockwise when cultures are illuminated from above and anticlockwise when illuminated from below. In *P. patens*, side branch initials (SBIs) emerge from caulonemal subapical cells on the outside of the curve. By contrast, the curvature of caulonemata of *Funaria hygrometrica* is predominantly anticlockwise when colonies are illuminated from above and clockwise when illuminated from below. In *F. hygrometrica*, SBIs emerge from caulonemal subapical cells on the inside of the curve. We have discounted a role for gravity in these phenomena and discuss several other possible mechanistic explanations. We also document for the first time thigmotropism of protonemata of *P. patens*.

Box 1.
Glossary of Terminology

| Box 1. Glossary of Terminology |  |
| --- | --- |
| Moss colony | A single plant derived from a spore or small inoculum |
| Protonema(-ta) | Filament(s) of tubular chloronemal or caulonemal cells |
| Chloronema(-ta) | Photosynthetic protonema(-ta) originating from spore germination, dedifferentiation of moss inoculum cells of several kinds, e.g. leaf cells, or, in the case of secondary chloronemata, from side branch initials |
| Caulonema(-ta) | Protonema(-ta) responsible for rapid colonisation of surrounding substratum and resulting from the differentiation of chloronemal apical cells or side branch initials into caulonemal apical cells |
| Side branch initial | A side branch comprised of a single cell formed by mitosis at the distal end of a caulonemal subapical cell |
| Gametophore | A photosynthetic bud or leafy shoot ultimately bearing gametes |
| Rhizoid | A tubular filament, usually but not exclusively, arising from the base of a gametophore and responsible for attachment to the substratum |
| Gravitropism | A growth response directed by gravity |
| Phototropism | A growth response directed by light |
| Thigmotropism | Redirected growth in response to a touch stimulus |
| Nutation | A bending movement of a plant organ caused by differences in growth in different parts of the organ |

## Introduction

A spiral organisation of protonemal filaments in a moss, *Dicranum*, was first described by Gause (1931). More detailed observations were made by Bopp (1959) mainly on *F. hygrometrica*. Relatively recently, Kern and co-authors described the occurrence of a spiral arrangement of protonemata of *Ceratodon purpureus* grown in darkness and microgravity during spaceflight (Kern et al. 2005) Preliminary observations of *P. patens* had shown that its caulonemal filaments exhibit predominantly clockwise curvature on the surface of solid growth medium when illuminated from above. Computer modelling suggested that this is important for the generation of a normal morphology, including the establishment of circular symmetry by protonemal colonies (Fracchia and Ashton 1995). Here we examine the curvature of *P. patens* and *F. hygromrtrica* caulonemata and its possible relationship to the formation of oblique cross walls by caulonemal apical cell division, hormonal communication between cells, nutation and tropisms.

## Materials and Methods

The *P. patens* line used in this study was derived from a subculture of Gransden 1962 *P. patens* obtained from a single spore isolated from nature by H.L.K. Whitehouse (Ashton and Cove 1977). *NicB5ylo6* was obtained by *N*-methyl-*N*′-nitro-*N*-nitrosoguanidine (NTG) mutagenesis of this wild-type line (Ashton and Cove 1977). The *F. hygrometrica* line was also derived from a single spore. Plants were grown axenically on solid ABC medium (Knight et al. 1988) supplemented with nicotinic acid (8 μM), with or without di-ammonium (+) tartrate (5 mM) under continuous white light (photon flux, 50-70 μmol m^-2^ s^-1^) at approx. 22-25 °C for 3 to 5 weeks. In some cases, the medium was overlaid with sterile, porous cellophane discs (Grimsley et al. 1977), in turn covered with a thin layer of agar medium (1-2 mm deep). The following lighting arrangements were employed.

1. Petri dishes were placed right side up on a white shelf, then covered with one layer of clear resin filter (Roscolux No. 114, Hamburg Frost) (to reduce the rate of evaporative water loss from the culture medium) and illuminated from above, i.e. through the lid of the dish.
2. Petri dishes were inverted, i.e. placed upside down on the shelf, covered with a layer of clear resin filter and illuminated from above, i.e. through the bottom of the dish and through the culture medium.
3. Petri dishes were placed right side up on a raised pane of clear glass, covered with a layer of black card (to prevent extraneous light impinging on the cultures from above) and illuminated from below, i.e. through the pane of glass, the bottom of the Petri dish and the culture medium.

Numerical data were recorded in Microsoft Excel files. Averages and sample standard deviations were then calculated using inbuilt formulae in the programme followed by rounding to two significant figures.

## Results

Caulonemal filaments of *P. patens*, when illuminated from above (lighting arrangement 1), grew over the surface of solid agar medium perpendicular to the incidence of light or slightly negatively phototropically into the medium. Those growing over the surface of the medium curved at a rate of approximately 1° per apical cell cycle (Table 1). The direction of curvature depended on the direction of the incident light. When right side up Petri dishes, which had been lit from above (lighting arrangement 1), were viewed through the Petri dish lid, the curvature was predominantly clockwise; when inverted, i.e. upside down, Petri dishes, which had been lit from above (lighting arrangement 2), were viewed through the Petri dish lid, it was predominantly anticlockwise (Table 2). Cultures initially illuminated from above (lighting arrangement 1) for 18 d and subsequently from below (lighting arrangement 3) for 9 d resulted in a reversal of the direction of curvature of caulonemata from clockwise to anticlockwise. By counting the number of new cells made with anticlockwise curvature and knowing how long it had taken to form them, we were able to estimate the cell cycle time of caulonemal apical cells, which had a duration of approximately 10 h (Table 3).

**Table 1.** Rate of curvature in degrees of *P. patens* caulonemal apical cells.

| Filament number | Number of cells per caulonemal filament | <sup>a</sup>Curvature of filament in degrees | Curvature in degrees per cell |
| --- | --- | --- | --- |
| 1 | 71 | 82 | 1.2 |
| 2 | 64 | 60 | 0.94 |
| 3 | 57 | 70 | 1.2 |
| 4 | 68 | 60 | 0.88 |
| 5 | 78 | 68 | 0.87 |
| <sup>b</sup>Ave | 68 | 68 | 1.0 |
| <sup>c</sup>StDev.S | 7.8 | 9.1 | 0.17 |

<sup>a</sup>Angle in degrees between a line through the central long axis of the caulonemal basal cell and a line through the central long axis of the caulonemal apical cell.
<sup>b</sup>Ave = average of values in each column (n = 5).
<sup>c</sup>StDev.S = Sample standard deviation of values in each column.
Values are given to two significant figures.

**Table 2.** Direction of curvature of *P. patens* and *F. hygrometrica* caulonemata.

| *P. patens* | Illuminated through Petri dish lid<br>(lighting arrangement 1) |  |  | Illuminated through bottom of Petri dish<br>(lighting arrangement 2) |  |  |
| --- | --- | --- | --- | --- | --- | --- |
|  | Clockwise | Straight | Anticlockwise | Clockwise | Straight | Anticlockwise |
| <sup>a</sup>Ave | **72** | 17 | 11 | 12 | 24 | **63** |
| <sup>b</sup>StDev.S | 17 | 14 | 14 | 8.7 | 7.3 | 6.8 |
| <sup>c</sup>n | 31 |  |  | 10 |  |  |
| *F. hygrometrica* | Illuminated through Petri dish lid<br>(lighting arrangement 1) |  |  |  |  |  |
|  | Clockwise | Straight | Anticlockwise |  |  |  |
| Ave | 2.4 | 4.5 | **93** |  |  |  |
| StDev.S | 3.3 | 3.6 | 6.0 |  |  |  |
| n | 5 |  |  |  |  |  |

<sup>a</sup>Ave = average of percentages of caulonemal filaments/gametophytic colony exhibiting clockwise, straight or anticlockwise curvature for n colonies. The highest values are shown in bold. Similar data from gametophytes grown on nitrate as sole nitrogen source and from colonies grown on nitrate plus ammonium were combined. All cultures were scored from the perspective of the Petri dish lid.
<sup>b</sup>StDev.S = Sample standard deviation of scores for n colonies.
<sup>c</sup>n = number of colonies.
Values are given to two significant figures.

**Table 3.**
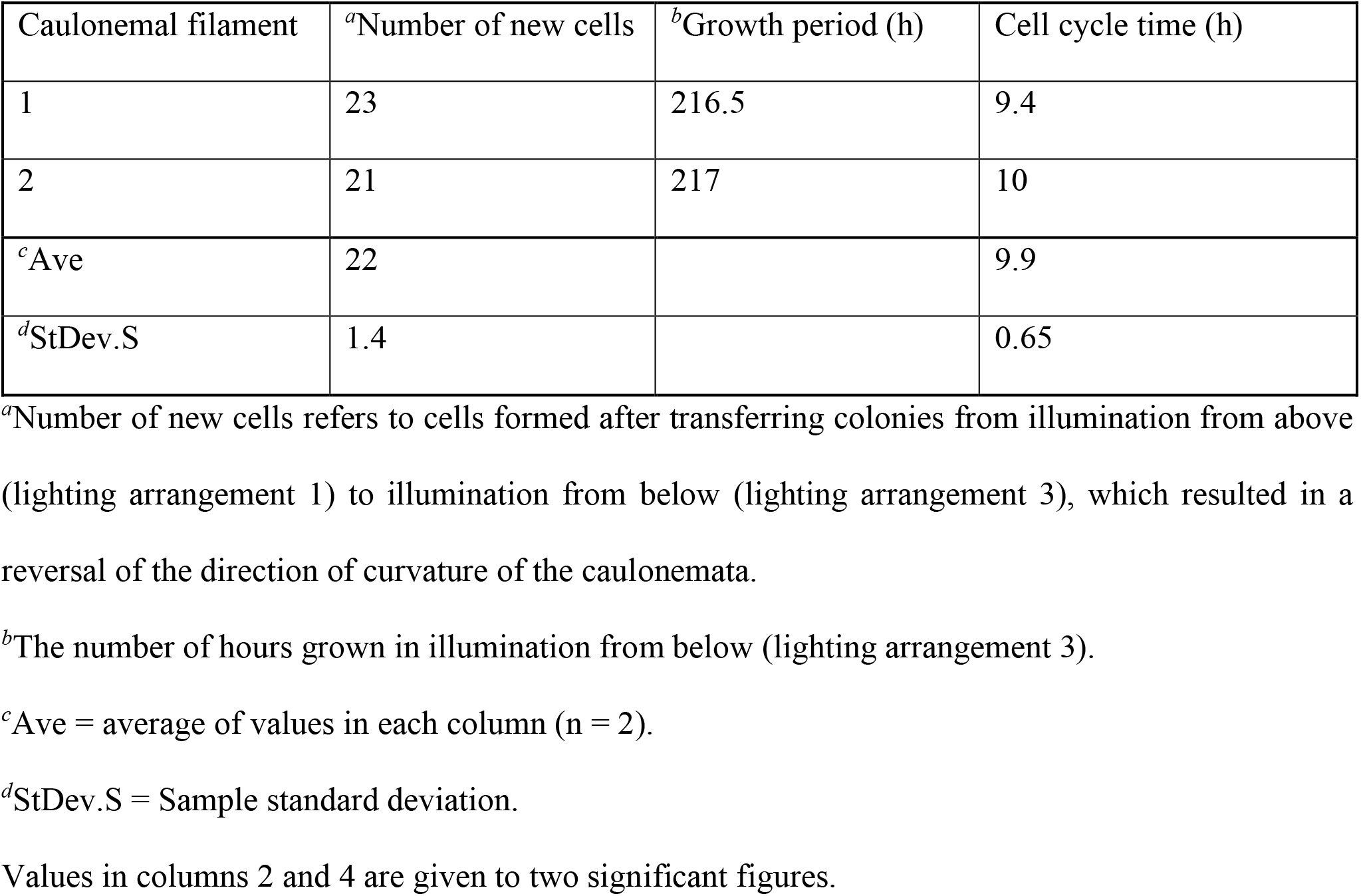
Caulonemal apical cell cycle time.

The direction of curvature of caulonemata of *F. hygrometrica*, when illuminated from above (lighting arrangement 1), was predominantly anticlockwise (Table 2).

Also of interest regarding the curvature of caulonemata was the sidedness of emergence of SBIs. SBIs emerged from the second or third subapical cell of caulonemata at the distal end of the cell and next to the acute angle generated by the oblique cross wall. In *P. patens*, the SBIs always emerged on the outside of the curve and developed predominantly into positively phototropic, branching secondary chloronemata or into secondary caulonemata or gametophores (Figure 1A and B). The consistent emergence of SBIs on the outside of the curve corresponded to the consistent repetitive orientation of oblique cross walls with respect to the long and short axes of the tubular caulonemal cells. Contrastingly, when illuminated from above, caulonemata of *F. hygrometrica* exhibited predominantly anticlockwise curvature (Table 2) with secondary protonemal branches emerging on the inside of the curve (Figure 1C).

**Fig 1A.**
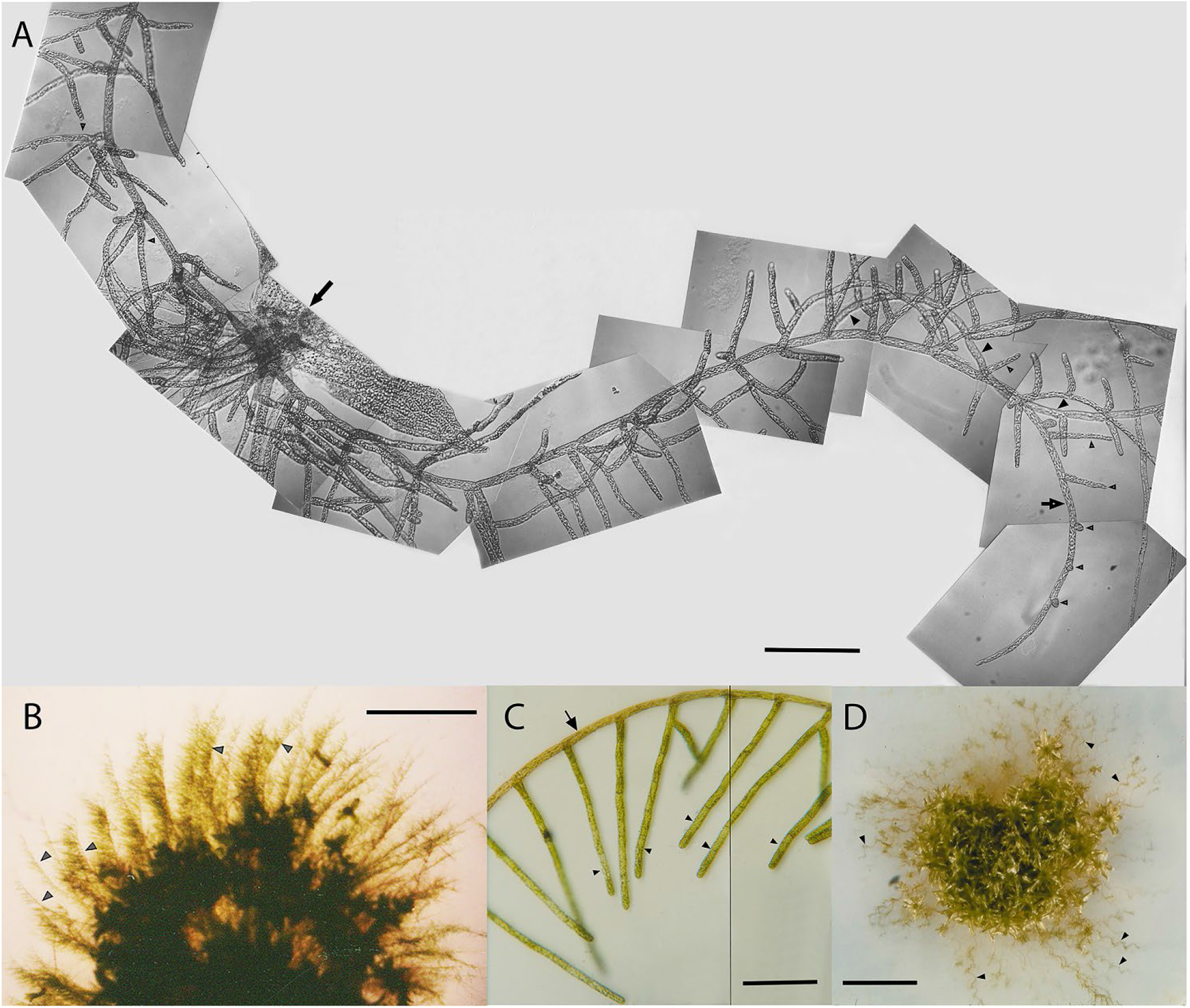
Composite image of an individual caulonema of *P. patens*. Short arrow points to a curved, primary caulonemal filament. Small arrowheads indicate SBIs, simple secondary chloronemal filaments and compound chloronemata emerging on the outside of the curves of the primary caulonema. Large arrowheads indicate a secondary caulonema, also emerging on the outside of the curve. Long arrow points to the base of a leafy gametophore with rhizoids. **B**. Half of a gametophytic colony of *P. patens* with spiral morphology. Arrowheads indicate peripheral caulonemata with clockwise curvature and secondary chloronemata emerging on the outside of the curve. **C**. Composite image of part of an individual caulonema of *F. hygrometrica*. Arrow points to a primary caulonema with anticlockwise curvature. Arrowheads indicate secondary chloronemata emerging on the inside of the curve. **D**. A gametophytic colony of *P. patens nicB5ylo6*. Arrowheads point to wavy, peripheral primary caulonemata. All cultures were illuminated in continuous white light from above (lighting arrangement 1). Scale bars represent 300 μm (**A, C)** and 5 mm (**B, D**).

We also discovered that, when caulonemal filaments were grown in a thin layer of agar medium, the crowded filaments contacted each other and wound around one another producing a multi-stranded, rope-like structure (Figure 2A and B). Individual caulonemata in the rope curved helically, while the multi-stranded ‘caulonemal rope’ was more or less straight.

**Fig 2A.**
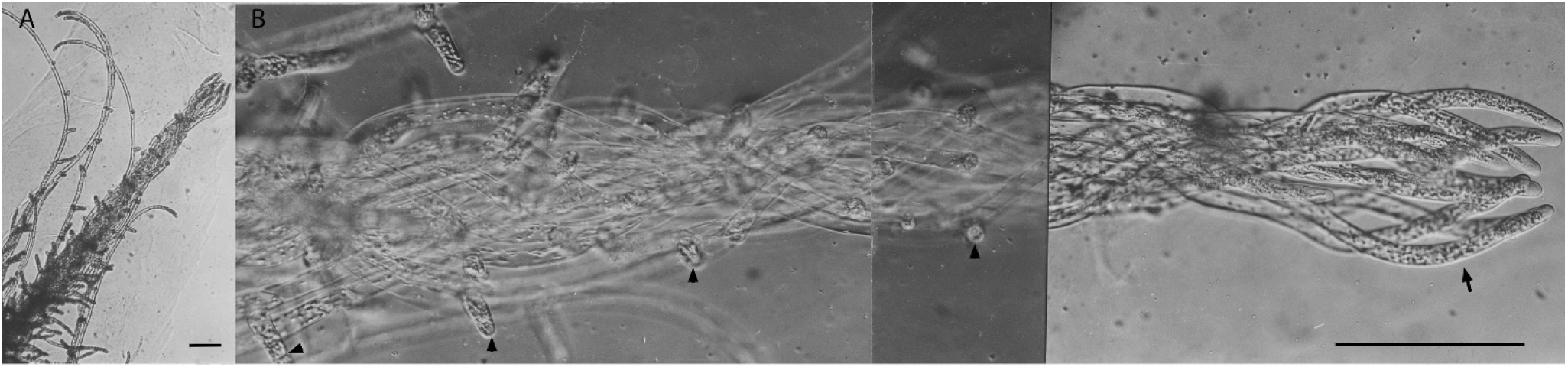
Four individual, curved caulonemata with side branch initials and secondary chloronemata and a ‘caulonemal rope’ comprised of multiple, intertwined caulonemata. **B**. Enlarged, composite image of a ‘caulonemal rope’. Arrow points to caulonemal apical cells; arrowheads indicate SBIs and developing secondary chloronemata. Scale bars represent 150 μm.

## Discussion

Our quantitative data on the curvature of *F. hygrometrica* caulonemata confirm Bopp’s qualitative observation that *F. hygrometrica* caulonemata curve predominantly anticlockwise when illuminated from above. Bopp also showed that the direction of curvature changes from anticlockwise to clockwise when cultures are illuminated from below (Bopp 1959). This contrasts with the behaviour of *P. patens* in which caulonemal curvature is predominantly clockwise when illuminated from above (lighting arrangement 1) and anticlockwise when illuminated from below (lighting arrangements 2 and 3). It is important to note that these statements are correct only when observations at the end of the growth period are made from the perspective of the Petri dish lids, e.g. through the lids, in all cases. When considered from the perspective of the light source, the orientation of curvature of *P. patens* caulonemata remains clockwise for all lighting arrangements. The same argument can be applied to Bopp’s data in which case curvature remains anticlockwise whether illuminated from above or below when scored from the perspective of the light source. Currently, we don’t have an explanation for these differences between two closely related members of the Funariaceae cultured in the same laboratory conditions. Conceivably, they may have relevance for these mosses when growing in their respective natural habitats.

Two other interesting questions about the phenomenon described above are:

1. What causes the curvature of caulonemata and the associated sidedness of the emergence of SBIs?
2. What selective advantage drove the evolution of these features?

Below we speculate on each question in turn and by a process of logical elimination attempt to reach conclusions. First, however, we think it is worth pointing out that, if we make the reasonable assumptions that primary caulonemal apical cells differentiate from centrally located, primary chloronemal apical cells at about the same time and radiate outwards with a constant cell cycle time, we can begin to understand why the population of peripheral caulonemata behaves in a highly coordinated manner. This will be because all the caulonemal apical cells will have approximately the same age and be approximately the same distance from the centre of the protonemal colony. It follows that these cells should and do respond to environmental challenges in a coordinated way that facilitates observation and interpretation of changes in colony morphology.

### Possible answers to question 1

#### Caulonemal curvature may result from an aberrant tropic response

As in the roots of *Arabidopsis* seedlings (Okada & Shimura, 1990), caulonemal curvature might be the result of an aberrant tropic response on the slippery agar surface. However, it is unclear whether the tropism concerned could be thigmotropic or phototropic.

It has been reported that rhizoids of some moss species, e.g. *Fontinalis squamosa* (Glime 1987) and *Calliergon stramineum* (Duckett 1994), are thigmotropic and coil tightly around any solid object (Goode et al. 1992). The main axes of rhizoids frequently wind around each other and fine ramifications of rhizoids wind around larger rhizoids or around caulonemata forming ‘rhizoid wicks’ or ‘rhizoidal ropes’ respectively (Duckett et al. 1998; Duckett et al. 2004). These are especially well developed in the Polytrichales (Wigglesworth 1947) and resemble quite closely the ‘caulonemal ropes’ we describe above. However, our report is of the first documented case of thigmotropism of protonemata of *P. patens* and appears to be limited to caulonemal filaments since secondary chloronemal filaments arising from caulonemata did not display this behaviour. We propose that the formation of ‘caulonemal ropes’ is a touch response, i.e. genuine thigmotropism, since it is unlikely that nutation could maintain the closely intertwined association of multiple caulonemata over the full length of a rope. Our discovery of caulonemal thigmotropism in *P. patens* raises the possibility that an aberrant thigmotropic response is responsible for the curvature of caulonemata on the slippery surface of solid agar medium.

A phototropic mechanism is supported by our observations that, at high photon flux as employed by us, *P. patens* caulonemata grow over the surface of solid agar medium perpendicular to the incidence of light or slightly negatively phototropically into the medium. This is the phototropic behaviour of caulonemata that typically occurs at high photon flux (Cove et al. 1978; Cove and Knight 1987). Also, as reported herein and by Bopp (1959), the direction of caulonemal curvature of both *P. patens* and *F. hygrometrica* appears to be set by the direction of incident light from the light source. These findings suggest that, as a mechanistic explanation of filament curvature, the occurrence of an aberrant phototropic response on the surface of slippery agar medium is equally as plausible as an aberrant thigmotropic response.

While we cannot discount the possibility that an aberrant tropism is the cause of caulonemal curvature at the solid agar surface or at least is a contributory factor, explanation of the occurrence of similarly curved caulonemata below the surface of the medium, which we have observed, is problematic. We believe gravity has no role in determining the direction of curvature in light-grown cultures for the following reasons. 1. Gravitropic responses in mosses are turned off by light at high photon fluxes such as those used in our study (Cove and Knight 1987). 2. Gravity is uniform across whole moss colonies in our experiments but, while the curvature of caulonemata was predominantly clockwise or anticlockwise depending on the direction of light, colonies were also characterised by the possession of collections of straight caulonemata or caulonemata of opposite curvature in regions of the colonies distinct from that displaying the predominant curvature (Table 2). Also, very occasionally the direction of curvature of an individual caulonemal filament was reversed (Figure 2A). We believe the most probable explanation for this is non-uniformity of the direction of light resulting from reflection from the bottom of the Petri dish.

#### Caulonemal curvature may be caused by apical cell nutation

Clockwise curvature of *Ceratodon purpureus* protonemata has been observed in space under conditions of microgravity and the absence of light by Kern et al. (2005). As is the case with moss caulonemata grown in the light, aberrant gravitropism cannot be the cause of this since the negative gravitropism of this moss, which occurs at ground level where gravity is 1 *g*, is absent in space. An alternative mechanistic explanation is that the protonemal apical cell of *C. purpureus* nutates in a circular or elliptical manner and slides on the surface of the slippery, solid agar medium. The direction of nutation, clockwise or anticlockwise, would then determine the direction in which the apical cell slides and thus the direction in which it curves. Assuming this is the same phenomenon we observed in *P. patens* and *F. hygrometrica* grown in the light, we can conclude that the direction of nutation is set differently in these two mosses by the direction of incident light from the light source. We can also infer that, if nutation involves rotation around the long axis of the caulonemal apical cell, it must be precisely synchronised with the formation of oblique dividing cross walls since otherwise the emergence of SBIs would be expected to occur on all sides of caulonemal subapical cells instead of only on the outside of the curve. Little has been written about nutation in bryophytes, but time-lapse photography appears to show small alternating changes of growth direction of elongating caulonemal apical cells of *F. hygrometrica* (Bopp and Brandes 1969). Also, we have obtained, by mutagenesis with NTG, a line of *P. patens, nicB5ylo6*, which is partly characterised by wavy caulonemal filaments (Figure 2D). This phenotype is very similar to the wavy phenotype of young seedlings of *Arabidopsis* growing on the surface of solid agar medium inclined at an angle of 45° to the direction of gravity (Okada and Shimura 1990) and we propose that in both cases it might be caused by repeated reversal of the direction of nutation and thus of sliding on the slippery agar medium. We also suggest that *nicB5ylo6* possesses one or more additional mutations responsible for the wavy phenotype since it seems unlikely that the *nic* or *ylo* mutation is the cause. It is interesting that at least six genes influence the wavy phenotype of *Arabidopsis* seedlings (Okada and Shimura 1990). Kern and coauthors (Kern et al. 2005) claimed that nutation couldn’t be the cause of protonemal curvature in *C. purpureus* in space since no nutations had been detected in dark-grown protonemata at 1 *g* (Young and Sack 1992). However, small bending deviations, like those discernible at the tips of light-grown *F. hygrometrica* caulonemal apical cells (Bopp and Brandes 1969), might easily be missed or alternatively completely masked by the very strong gravitropic response of *C. purpureus* protonemata grown in the dark at ground level. Therefore, we suggest that aberrant nutation remains a viable explanation of the spiral morphology of light-grown moss and of moss grown in microgravity and darkness.

#### Caulonemal curvature, and the sidedness of emergence of SBIs, in light may be directed by consistent orientation of oblique cross walls formed by division of caulonemal apical cells

Although we do not know whether the consistent orientation of oblique cross walls plays a causative role in filament curvature or is a consequence of it or is unrelated to it, it is plausible that curvature of the apical cell generates asymmetrical tensions in the cytoskeleton that in turn could be responsible for repositioning the mitotic spindle and cell plate during cell division resulting in the formation of an oblique cross wall. Evidence supporting this was obtained by Bopp and Brandes (1969), who used time-lapse photography of *F. hygrometrica* to demonstrate that the spindle and cell plate were re-orientated respectively from being in line with and perpendicular to the central long axis of the caulonemal apical cell to positions corresponding to the oblique cross wall subsequently laid down. Additional circumstantial evidence supporting this mechanism is that, whereas curved caulonemata possess oblique dividing cross walls, phototropically responsive chloronemata have transverse cross walls, which are perpendicular to the long axis of the cells, and lack curvature after their growth has been reoriented by light, growing directly towards light in the case of secondary chloronemata and approximately at a right angle to it in the case of primary chloronemata.

As noted earlier (see Results), SBIs emerge from the second or third subapical cell of caulonemata at the distal end of the cell and next to the acute angle generated by the oblique cross wall. Thus, as is the case with curvature, there is a very strong relationship between the sidedness of emergence of SBIs and the consistent orientation of caulonemal cross walls.

#### Cell to cell communication via a diffusible inhibitor may be responsible for spiral morphology

In 1959, Bopp reported that a diffusible factor made by *F. hygrometrica* inhibits caulonemal elongation and the expansion of gametophytic colonies. He subsequently named this substance Factor H and showed that it originates from the caulonemata themselves (Bopp 1963). More recently, Proust and coauthors reported that, in 3-week old *P*.*patens*, strigolactones originating from gametophores act as diffusible signals inhibiting caulonemal SBI formation, caulonemal elongation and colony expansion in a colony density-dependent way (Proust et al. 2011). Based on differences in their chemical properties and in the timing of their morphological effects, Factor H is probably not a strigolactone. It is also unlikely that strigolactones are the cause of spiral morphology, which is already established by the time strigolactones have accumulated to sufficiently high levels to inhibit the elongation of caulonemata. Nevertheless, the possibility that a diffusible inhibitor communicates between caulonemal apical cells coordinating their behaviour and keeping them separated remains attractive. The acquisition of spiral morphology in which caulonemata are curved, predominantly in the same direction, and relatively evenly spaced apart is more difficult to understand but might be achieved if the responsible diffusible inhibitor is only effective over a short distance.

We would prefer to conclude that a single mechanism underlies the acquisition of a spiral morphology by moss protonemal colonies under all circumstances. However, all our observations and those reported by others have been made in precisely controlled laboratory conditions that may not be replicated exactly in nature. Therefore, some mechanisms, e.g. an aberrant tropism on a slippery surface, may make a contribution in the laboratory but have no relevance in the moss’s natural habitat. Also, we can discount a role for gravity in the light and in the dark in microgravity. Therefore, if we must propose based on current information an explanation that applies to all situations, we suggest that spiral morphology of protonemal colonies is caused by a combination of nutation and the formation of consistently oriented oblique cross walls by caulonemal apical cells whose behaviour may be at least partially coordinated by a short-acting diffusible inhibitor.

### Possible answers to question 2

Spiral arrangements of plant organs, e.g. the leaves of *P. patens* and *F. hygrometrica* gametophores, are common and have presumably evolved to minimise overlap between them thereby maximising various processes including and notably photosynthesis. The spiral organisation of caulonemata coupled with the emergence of photosynthetic secondary chloronemata and leafy gametophores on one side of them could be another example of this. Also, caulonemata are the organs mainly responsible for rapid colonisation of the surrounding substratum. Our estimate of the caulonemal cell cycle, approximately 10 h, accords fairly well with that, approximately 7-8 h, reported by other researchers (Cove 1992; Vidali and Bezanilla 2012) and is about two to three times faster than the cell cycle time of chloronemal apical cells, approximately 20-24 h (Cove 1992; Vidali and Bezanilla 2012). It is easy to imagine how the emergence of secondary caulonemata on the outside of curved primary caulonemata of *P. patens* might lead to a spreading morphology further promoting colonisation of new areas containing previously untapped sources of nutrients such as mud and seasonally moist soil at the edges of ponds, one of its natural habitats. By contrast, the emergence of secondary branches on the inside of curved caulonemata of *F. hygromeyrica* should favour the production of a more compact morphology, which we assume provides some ecological benefit in this moss’s natural habitat. Finally, if spiral protonemal morphology evolved to optimise photosynthesis and if this phenomenon is the same as that overridden by gravitropism in the dark, there would probably have been insufficient selective advantage for its evolutionary loss.

## Notes

### Competing Interest Statement

The authors have declared no competing interest.

### Summary of Updates

Discussion has been revised and expanded.

